# Genomics via Optical Mapping (I): 0-1 Laws for Mapping with Single Molecules

**DOI:** 10.1101/000844

**Authors:** Thomas Anantharaman, Bud Mishra

**Affiliations:** BioNano Genomics, San Diego, CA 92121.; Courant Institute of Mathematical Sciences, New York University, 251 Mercer Street, New York, NY, USA 10012.; Cold Spring Harbor Lab, 1 Bungtown Road, Cold Spring Harbor, NY, USA 11724.

## Abstract

The genomic data that can be collected from a single DNA molecule by the best chemical and optical methods (e.g., using technologies from OpGen, BioNanoGenomics, NABSys, PacBio, etc.) are badly corrupted by many poorly understood noise processes. Thus, single molecule technology derives its utility through powerful probabilistic modeling, which can provide precise lower and upper bounds on various experimental parameters to create the correct map or validate sequence assembly. As an example, this analysis shows how as the number of “imaged” single molecules (i.e., coverage) is increased in the optical mapping data, the probability of successful computation of the map jumps from 0 to 1 for fairly small number of molecules.

## 1 Some Preliminary Remarks

Optical Mapping [AMS97, Ana+97b, AnMS99, AsMS99, Cai+98, Mishra03, Sam+95] is an approach that generates an ordered restriction map of a DNA molecule (e.g., a genome [Jing+99, Lai+99, Lim+01, lin+99, Zhou+02] or a clone [Cai+98, Giacalone+00, Sam+95, Skiadas+99]). The resulting restriction map is represented as an ordered enumeration of the restriction sites along with the estimated lengths of the restriction fragments between consecutive restriction sites and various related statistics. These statistics accurately model the errors in estimating the restriction fragment lengths as well as the errors due to unrepresented and misrepresented restriction sites in the map. These physical maps have found applications in improving the accuracy and algorithmic efficiency of sequence assembly, validating assembled sequences, characterizing gaps in the assembly and identifying candidates for finishing steps in a sequencing project. Also, because of its inherent simplicity and scalability as well as its reliance on single molecules, optical mapping also provides a fast method for moderate resolution karyotyping and haplotyping.

The physico-chemical approach underlying optical mapping is based on immobilizing long single DNA molecules on an open glass surface, digesting the molecules on the surface and visualizing the gaps created by restriction activities using fluorescence microscopy. Thus the resulting image, in the absence of any error, would produce an ordered sequence of restriction fragments, whose masses can be measured via relative fluorescence intensity and interpreted as fragment lengths in base pairs. The corrupting effects of many independent sources of errors affect the accuracy of an optical map created from one single DNA molecule, and can *only* be tamed by combining the optical maps of many single molecules covering completely or partially the same genomic region and by incorporating accurate statistical models of the error sources. To a rough approximation, the insurmountable obstacles in the chemistry is circumvented by cleverly exploiting the statistical properties of the system through a “0-1 Law” in the parameter space. This law plays a crucial role at the heart of the entire optical mapping technology and is likely to reappear in other contexts as well, e.g., array-mapping, single-molecule sequencing and haplotyping. In this paper, we focus on one such law in the context of mapping clones and we hint at how these results are generalized to genomic mapping.

The main error sources limiting the accuracy of an optical map are due to either incorrect identification of restriction sites or incorrect estimation of the restriction fragment lengths. Since these error sources interact in a complex manner and involve resolution of the microscopy, imaging and illumination systems, surface conditions, image processing algorithm, digestion rate of the restriction enzyme and intensity distribution along the DNA molecule, statistical Bayesian approaches are used to construct a consensus map from large number of imperfect maps of single molecules. In the Bayesian approach, the main ingredients are as follows: (1) A model of the map of restriction sites (*Hypothesis*, *H*) and (2) A conditional probability distribution function for the single molecule map data given the hypothesis (*Conditional pdf*, *f* (*D*|*H*)). The conditional pdf models the restriction fragment sizing error in terms of a Gaussian distribution, the missing restriction site event (due to partial digestion) as a Bernoulli trial and the appearance of false restriction sites as a Poisson process. Using the Bayes’ formula, the posterior conditional pdf 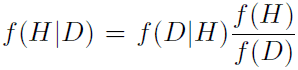 is computed and provides the means for searching for the best hypothetical model given the set of single molecule experimental data. Since the underlying hypothesis space is high dimensional and the distributions are multi-modal, a natïve computational search must be avoided. An efficient implementation involves approximating the modes of the posterior distribution of the parameters and accurate local search implemented using dynamic programming [AMS97, AnMS99]. The correctness of the constructed map depends crucially on the choice of the experimental parameters (e.g., sizing error, digestion rate, number of molecules). Thus, the feasibility of the entire method can be ensured only by a proper experimental design.

This paper studies several simple models for optical mapping and explores their power and limitations when applied to the construction of maps of clones (e.g., lambdas, cosmids, BACs and YACs), by providing precise lower and upper bounds on the number of clone molecules needed to create the correct map of the clone. Our probabilistic analysis proves the existence of a 0-1 laws in the number of molecules.

The paper is organized as follows: In section 2, we formulate the problem; in sections 3, 4 and 5, we successively introduce and analyze the effects of various error sources: namely, partial digestion error, misorientation error and quantization error, respectively. We use probabilistic methods to provide upper and lower bounds on the choices of parameters that would ensure correct result with high probability. In section 6, we study the effect of sizing error and its interaction with discretization. The analysis indicates that for a reasonable choice of sizing error, the algorithms based on discretization are unlikely to work correctly with any reasonable probability.

## 2 Problem Formulation

The underlying bio-chemical problem concerns with the construction of an ordered restriction map of a clone (a piece of DNA of length *L*, where *L* is measured in base-pairs *bp*s, *Kb* = 10^3^*bp* or *Mb* = 10^6^*bp*). Typical values of *L* are 2-20*Kb* (lambda's), 20-45*Kb* (cosmids) 150–200*Kb* (BAC'S) and ≈ 1*Mb* (YAC’S). For our mathematical analysis, we will often assume that *L* takes some fixed value which can be arbitrarily large. These clones are sequences of length *L* over the alphabet { A, T, C, G }. Certain short subsequences (typically of length 6, e.g., GGATCC) can be recognized by a restriction enzyme (e.g., *BamH* I), and location of these restriction sites

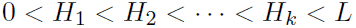

in the clone is the *ordered restriction map* of the clone with respect to the given enzyme.

Let *h_i_* = *H*_*i*_/*L* be a real number. Then the *normalized ordered restriction map* of the clone with respect to the enzyme is

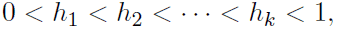

where each *h_i_* assumes some real value in the open unit interval (0, 1).

Note that in the absence of any additional distinguishing characteristic of the clone (e.g., identification of 3' end or 5' end), we could have also taken the following as another *normalized ordered restriction map* of the same clone with respect to the same enzyme:

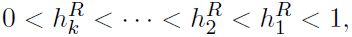

where 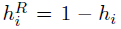. Note that the *normalized ordered restriction map* is unique up to reversal in the absence of any additional distinguishing characteristic, and is unique if we know the orientation.

## 3 False Negative Errors: Partial Digestion

Let us postulate an experiment, where the desired *normalized ordered restriction map* is observed, subject to *partial digestion* error and where any particular restriction site is observed with some probability *p* ≤ 1. We assume no other error sources for now; thus no other spurious sites (false restriction cuts) are included in the observation and the observed restriction map appears in the correct orientation.

Thus the result of the experiment is an ordered sequence of sites (normalized)

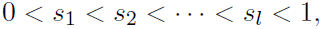

where for each *s_i_*, there is an *h_j_* in the true map, such that *s_i_* = *h_j_*. By assumption, for each *h_j_* the probability

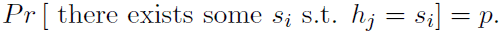

Let us also assume that the experiment is repeated *n* times resulting in *n* observed restriction maps. Assume that the true restriction map is unknown and is to be constructed from these *n* observations. A straight forward algorithm for doing this would be to simply take the union of all the observed restriction sites, and output this result in sorted order.

We claim that if 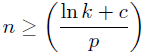 then the result of the preceding algorithm is almost surely correct. Here *c* is a constant to be determined later. Note that the probability that a cut site *h_j_* appears in at least one observation is 1 − (1 − *p*)^*n*^ ≥ 1 − *e*^−c^*/*k**. Thus the probability that all *k* true cut sites show up in the final map is given by (1 − *e*^−c^/*k*)^*k*^ > *e*^−*e*^−*c*^^ + *o*(1).

On the other hand, if 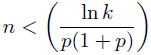 (*k* ≥ 1 and 0 < *p* < 0.69) then there is a high probability that the amount of data is insufficient to recover the correct map. Note again that given a true cut site *h_j_* the probability that this cut is never observed in any of the *n* observations is simply (1 − *p*)^*n*^ > *e*^−*pn*(1+*p*)^ > *e*^−ln*k*^ = 1/*k*. Thus, with this value of *n*, the probability that we can recover all the true cut sites is simply bounded from above by 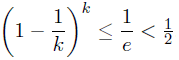.

In summary: Let *ϵ* be a positive constant and *c* ≥ ln(1/*ϵ*). Then for 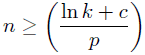 with probability at least (1 − *ϵ*), the correct ordered restriction map can be computed in *O(nk)* time.

When 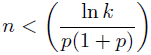 (*k* ≥ 1 and 0 < *p* < 0.69), there is a probability greater than half that no algorithm can compute the correct ordered restriction map.

*Note*, however, that since the value of *k* and *p* are not known a priori, it is impossible to use this result in a meaningful way in designing an experiment (i.e., in choosing *n*). The algorithm itself does not use the parameters *k* or *p*; only its success probability is determined by these parameters for a fixed set of input data.

## 4 Misorientation Errors

Next, let us postulate a modified experiment, where the desired *normalized ordered restriction map* is observed, subject to *partial digestion* error as well as error due to *mis-orientation*. Thus the result of the experiment is an ordered sequence of sites

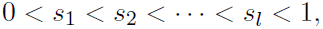

where either the sequence or its reversal

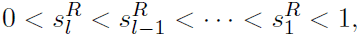

could be assumed to be derived from the true normalized ordered restriction map

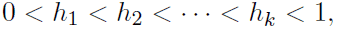

after partial digestion. By assumption, for each *h_j_* and for each observation, the probability

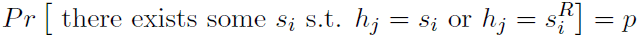

models the partial digestion.

**Assumption:**For the time being, we assume that the true normalized ordered restriction map has no *symmetric site*, i.e.,

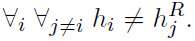

Let us also assume that the experiment is repeated *n* times resulting in *n* observed restriction maps whose orientations may be misspecified.

An algorithm to reconstruct the true map may proceed in two phases: In the first phase, all the molecules are folded by the mid-point, thus creating “folded-maps,” in which the orientation of the molecule is no longer an issue. The individual “folded-maps” are combined to compute a “consensus folded-map.” In the second phase, the “consensus folded-map” is unfolded back to create the restriction map where each individual site is assigned to either left half or right half, by examining the relative locations of pairs of restriction sites found in the original data (assuming that enough such information is available).

### 4.1 Phase 1

Define a function

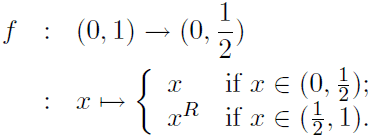

In phase 1, our goal is to construct the set

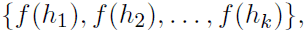

which can be easily accomplished by considering the sets

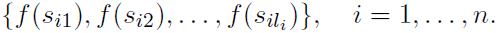

and proceeding in a manner similar to the one outlined in the preceding section. Using the arguments given earlier, we see that we will succeed in this phase with probability 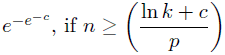.

### 4.2 Phase 2

While one cannot recreate the map directly from the result of the phase 1, one can invert *f* correctly, if each computed site is further augmented with a sign value (∈ {+1,−1}), where +1 denotes that the site belongs to the left half [(0, 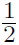)] and −1 denotes that the site belongs to the right half [(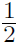, 1)]. Thus, we may define

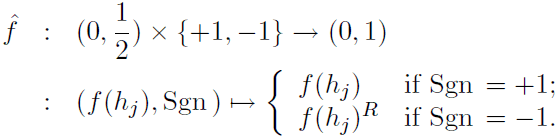

We can assign the sign values correctly as follows: Define a graph *G* = (*V*, *E*), where *V* = {*f*(*h*_1_), *f*(*h*_2_), …, *f*(*h_k_*)} and *e* = [*f*(*h_a_*), *f*(*h_b_*)] ∈ *E* if and only if

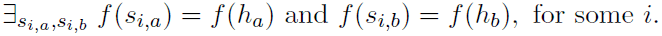

Furthermore, label *e* with +1 if for some *i*, either *s_i,a_* and *s_i,b_* ∈ (0, 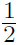) or *s_i,a_* and *s_i,b_* ∈ (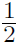, 1) (*both sites belong to the same half*); and with −1 if for some *i*, either *s_i,a_* ∈ (0, 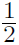) and *s_i,b_* ∈ (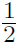, 1) or *s_i,a_* ∈ (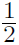, 1) and *s_i,b_* ∈ (0,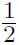) (*two sites belong to different halves*). In other words,

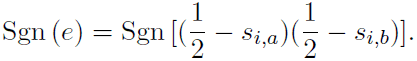

It is trivial to see that if the graph is *connected* then one can compute the correct vertex labels by first labeling an arbitrary vertex +1 (say, *f* (*h_j_*)) and then labeling the remaining vertices by following the edge labels during a graph-search process. Thus if *f* (*h_i_*) and *f* (*h_j_*) are path connected by a simple path *e*_1_, *e*_2_, …, *e*_m_ then>

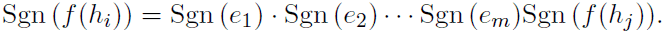

Next we compute an upper bound on the number of observations necessary for *G* to be connected. Let

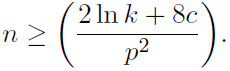

Let *S_h_*_1_ denote the set of observations with a cut site matching *f* (*h*_1_). The number of such observations, |*S_h_*_1_|, follows a Binomial distribution ∼ *S*(*n*, *p*).

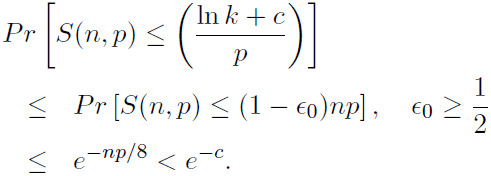

A cut corresponding to every *f*(*h_i_*) [2 ≤ *i* ≤ *k*] occurs in an observation in *S*_*h*1_, when 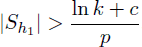, with probability

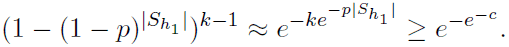

Thus *G* is connected with a probability > (1 − *e*^−*c*^)*e*^−*e*−*c*^ as [*f*(*h*_1_), *f*(*h_i_*)] appears in *G* for all 2 ≤ *i* ≤ *k*.

Summarizing: Let *ϵ* be a positive constant and *c* ≥ ln(2/*ϵ*). Then for

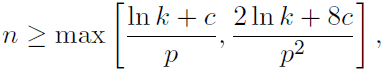

with a probability at least (1 − *ϵ*), the correct ordered restriction map can be computed in *O*(*nk*^2^) time. Also, see Appendix A1, for a slightly better bound when *p* = *O*(1/*k*).

### 4.3 Optical Cuts

Next we shall consider the situation where we have additional spurious cuts (optical cuts) that do not correspond to any restriction sites. A sound probabilistic model for these spurious cuts can be given in terms of a Poisson process with parameters λ_*f*_ (thus the expected number of false cuts per molecule is λ_*f*_). Hence, for any small region [*x*, *x* + δ*x*] in an observation,

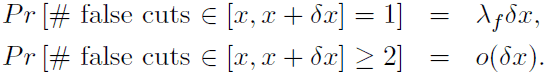

The probability that an observation contains exactly *f* spurious cuts is given by: 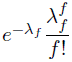. Typical observed values for λ*f* are about 0.2 for Lambda clones, 0.5 for cosmids and 1.0 for BAC’s. Thus, we expect roughly 1 false cut per 100*Kb*.

Under this model, it is fairly trivial to see that the false cuts pose no serious problem. Our algorithm can be modified in a straight forward manner where Phase 1 computation needs to be somewhat more robust.

In phase 1, our goal is to construct the set

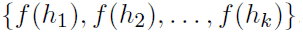

This is accomplished by considering the observation-based sets

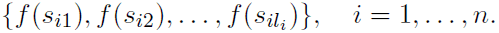

and including only those *f*(*s_ij_*)’s that occur at least twice in the combined observations. In other words, if there exists an *i*_1_ ≠ *i*_2_ such that if

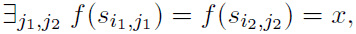

then include *x* in the output set.

Assume that 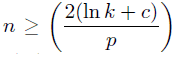, (with *c* > 1.26). Then if *h_i_* is a true cut site, the probability that *f*(*h_i_*) is not included in the output is

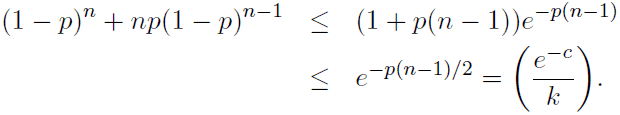

Proceeding as before the probability that all *k* true sites will be included is thus bounded from below by *e*^−*e*^−*c*^^, Also, by the assumption regarding the distribution of spurious cuts, we see that the probability that a spurious cut is included in the final set is zero.

### 4.4 Symmetric Cuts

Next, assume that the true ordered restriction map consists of *k* asymmetric cuts and *m* symmetric cuts. Thus the total number of cuts is *k* + 2*m*. Note that a cut *h_i_* is a symmetric cut, if both 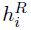 are true cuts. Additionally, we assume that the observations are subject to the *partial digestion* errors, *misorientation* errors, *spurious cut* errors (determined by a Poisson process) and *symmetric cuts*.

In this case, we proceed as before with phase 1 from the preceding subsection, and again assuming that 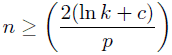, we will almost surely (with probability no smaller than 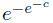) construct a set

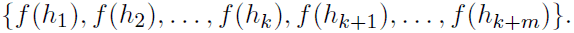

Note that the 2*m* symmetric sites yield *m* values in the folded structure when *f* is applied.

However, before proceeding to phase 2, we will remove those *f*(*h*_*j*_)‗s from the preceding set that correspond to symmetric cuts. A simple approach we can take is to check each observation for the existence of symmetric cuts at positions *s* and *s^R^*, where *f*(*s*) = *f*(*s*^*R*^) = *f*(*h_j_*).

We claim that if 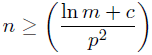 then the preceding steps correctly detect the symmetric cuts with probability greater than 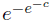. Note that assuming *h_j_* to be a symmetric true cut, the probability that the above test fails in any particular observation is (1 − *p*^2^) and thus the probability that the symmetric cut *h*_*j*_ goes undetected in any of the *n* independent observations is (1 − *p*^2^)^*n*^. Thus the probability that all *m* symmetric true cut sites are detected in the final map is given by [1 − (1 − *p*^2^)^*n*^]^*m*^ > 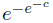.

Again, by the assumption regarding the distribution of spurious cuts, we see that the probability that a spurious cut is included or symmetric cut is missed in the final set is zero.

At the end of this step, we are left with a set containing only asymmetric cuts

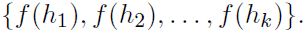

At this point, we simply proceed with the phase 2 *mutatis mutandis* and claim results similar to the ones derived earlier.

### 4.5 Summary

Consider an ordered restriction map with *k* + 2*m* restriction sites, of which *m* are symmetric cuts. Assume that the postulated experiment observes these maps, with each observation suffering from partial digestion error (*p* ≤ 1), misorientation error, spurious cuts (determined by a Poisson process with parameter λ_*f*_), but no sizing error.

**Theorem 4.1** *Let *ϵ* be a positive constant and c* ≥ ln(5/*ϵ*). *Then for*

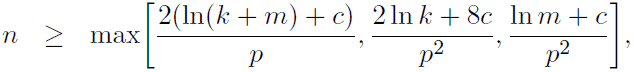

*there is a probability of at least (*1 − **ϵ*) that the correct ordered restriction map can be computed in O*(*n*(*L* + *k*^2^ + *m*)) *time*.

*When*

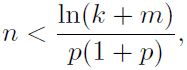

(0 < *p* < 0.69), *there is a probability greater than half that no algorithm can compute the correct ordered restriction map.*

## 5 Discretization

Next, we consider the effect of discretizing the map data by dividing it uniformly into several intervals of equal sizes. The main argument in favor of discretizing the data has been to accommodate the sizing errors that make the cut locations deviate from their true location. The main source for the sizing error has been the nonuniform attachment of the flurochromes that are necessary to visualize the DNA. We will study the effect of sizing error on discretization in a later section.

Let us assume that the clone DNA that we wish to analyze is of length *L* bps. Let ∆ represent a small subinterval and *δ* = ∆/*L*. Thus the unit length is partitioned into *M* = 1/δ = *L*/∆ consecutive subintervals. One assumes that it is not possible to distinguish the restriction cuts and spurious cuts in each of these subintervals. Thus, we need to ensure that *δ* is significantly small so that no more than one true restriction cut location belongs to a subinterval. We now write *r* = λ_*f*_ *δ* = ∆λ_*f*_/*L* to denote the probability that we shall observe one spurious cut in a subinterval. Note that the probability that we shall observe *f* spurious cuts in any observation is given by

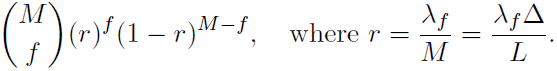

Thus in the limit as *M* → ∞ and *r* → 0,

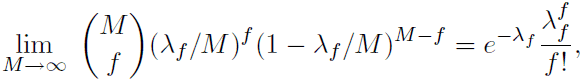

the analysis given earlier holds true. Furthermore, if we are simply interested in the effect of finite *M* (and nonzero *r*), we are still able to prove that for realistic values of *r* < *p*/27 the earlier bounds still hold. Thus, it suffices to ensure that λ*_f_*/*M* < *p*/27, or *M* > 27λ*_f_*/*p*—for instance, *M* could be 270 and satisfy the inequality as long as λ_*f*_ < 1.0 and *p* > 0.1. (See appendix A2.)

Typical values for various clones may be as follows: for lambdas, *M* can range from 200 to 2,000 and *r* ≈ 10^−3^–10^−4;^ for cosmids, *M* is 2,000–4,000 and *r* ≈ 10^−4;^ for BACs *M* ≈ 15,000 and *r* ≈ 10^−4^. In general, even for significantly smaller (but still realistic) values of *M*, *r* ≪ *p*.

### 5.1 Limit on *M*

It is worth noting that the discretization process makes it possible for spurious cuts to introduce a “wrong” cut site into final map. For instance, if each of the *n* observations contains a spurious cut in the same subinterval, then no algorithm can distinguish this spurious cut from a true cut (independent of digestion rate). Thus the probability that none of the *M* subintervals has a spurious cut in each of the *n* observations is given by

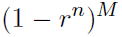

Now if we assume that 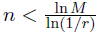, then the above probability is bounded from above by

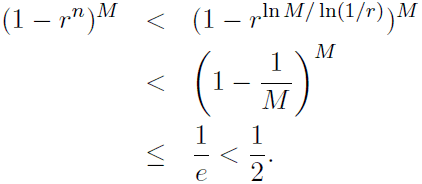

Hence we must further guarantee that

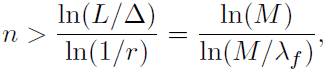

since otherwise there is a probability of half or more that the computed map will be wrong.

## 6 Sizing Errors

Next, suppose we model the sizing error and analyze its effect. Before doing so, we need to derive some inequalities relating the size of the discretized subinterval (∆) to several other external parameters. In particular, in order to infer the map correctly with probability greater than 1/√2, we must guarantee that 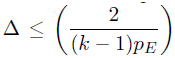, where *k* is the number of cuts and *p_E_* denotes the probability that the restriction enzyme cuts at a site.

Assume that 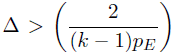. Let *l* denote the length of the smallest restriction fragment (piece of the molecule between two consecutive restriction sites). Note that the fragment lengths are distributed as 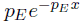, and the probability that a fragment is of length ≥ ∆/3 is

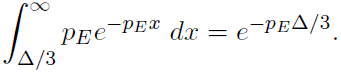

Thus the probability that the smallest of all (*k* − 1) fragments is no smaller than ∆/3, is

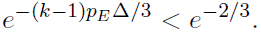

Thus the probability that the smallest fragment is of length ≤ ∆/3 and that both ends of the fragment belong to the same subinterval is bounded by

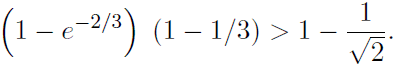

However, note that for a BAC clone, this implies that the largest value we may choose for ∆ ≤ 200*bp* (requiring *M* to be about 750).

Next assume that a true cut site at location *h* actually appears as a Gaussian distribution ∼ *N*(*h*, *σ*). Again, considering the complementary requirement to the one mentioned earlier, we must ensure that the observed cuts corresponding to the same true cut (at location *h_i_*) belong to the same subinterval with high probability. As a result, we may require that

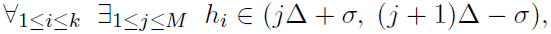

with high probability (say, ≥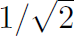). Thus, we require that

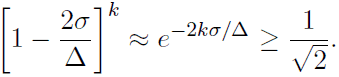

In other words, we require that 2*kσ*/∆ ≤ ln 2/2, and

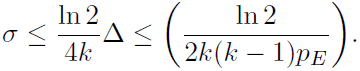

A simple calculation for the BAC example reveals that in order to guarantee the above inequality we need that *σ* ≤ 0.89*bp*. Thus for all practical purposes, in order for the discretized algorithm to work with any degree of correctness, we must require the *observation to be free of sizing error*. As a result, one can explain why several algorithms devised to work with dicretization failed, while purely continuous versions (or some combination) have done well.

## 7 Experimental Verification

This section compares the performance of a program based on the maximum likelihood approach to map-computation (described in [AMS97]) with the theoretical bounds in the previous sections. At the time of this writing, AMS algorithm [AMS97] still remains the *only* algorithm that has worked successfully on raw experimental data, without access to any extraneous parameters or the final answer. In each case, when the computed map was verified with data (from sequence and gel data) derived independently and subsequent to the experiment, the algorithm was found to be remarkably successful.

For all the experiments described in this section, random data were generated using the data models of the previous sections. For each data model and assumed number of data molecules, we generated 20 random data samples and counted the fraction of these samples for which the maximum likelihood program computed the correct map. For each data model the number of data molecules is varied to obtain the fraction of cases solved correctly as a function of the number of data molecules. We show that in each case there is a fairly sharp transition from not being able to solve any of the 20 samples to being able to solve all 20 samples. Moreover this transition point lies within the theoretical bounds computed in the previous sections. Finally we examine the performance of the maximum likelihood program for the case where there is significant sizing error. In this case the discrete methods described previously fail to work altogether, whereas the maximum likelihood method continues to work, albeit requiring a larger number of data molecules as the sizing error increases.

The maximum likelihood approach described in [AMS97] is based on a continuous (non-discrete) modeling of the data. The modeling of sizing error in the model results in a singularity in the probability density when the sizing error is zero. Therefore this case was approximated by assuming a small sizing error of 10^−5^ of the total molecule size, *σ* = 1.5bp. Each data model is specified by providing the number (*k*) and value of the actual cut locations, the sizing error in the form of a standard deviation (*σ*), a digest rate (*p*) and a false cut rate (λ_*f*_). For each model, random data is generated with the
help of a random number generator in a straight forward fashion: For each of the actual cuts, we draw a random number uniformly from [0,1], and if this value is below *p*, the cut is assumed to be present. We then draw another random number from the standard Gaussian distribution to determine the location of the cut, thus modeling the effect of sizing error. Next, false cuts are added by first drawing a random sample from a Poisson distribution with mean λ*_f_* to determine the number of false cuts, and then drawing the required number of random samples uniformly over [0,1] to get the false cut locations.

This step results in the generation of one *in-silico* “molecule.” This process is repeated to get the required number of molecules to make up one data set. This data set is then input as raw data to our maximum likelihood program, and the resulting map is scored a success if the number of cuts is correct, and the location of each cut is within one standard deviation (*σ*) of the correct location. (Note that *σ* is the standard deviation for the cuts of one sample molecule: the map computed by the AMS algorithm typically has a sizing error much less than that since the data from all molecules are averaged). This process is repeated for a total of 20 samples and the fraction of times the program succeeds is recorded against the data sample parameters (*k*, *σ*, *p*, λ*f*, number of molecules). The whole process (i.e., the one generating 20 samples) was repeated for different values of the parameters. The number of cuts was varied using the values *k* = 0, 1, 2, 5, 10, 20 and 37. The values of *p* tested were *p*=0.10 and *p*=0.20. The values of λ_*f*_ tested were λ_*f*_=0 (no false cuts) and λ_*f*_=1,2 and 4. For most experiments we selected *σ* = 1.5bp to approximate no sizing error, but for a small number of experiments with *k* = 2 and 37 we also tested *σ* = 150bp, 300bp, 750bp and 1.5Kb. Most experiments were repeated with the number of molecules set at 10, 20, 30, 40, 50, 70, 100, 200, 500 and 1000, and in a few instances 2000 or 5000.

The results are summarized in a series of graphs showing the success rate (out of 20 samples) as a function of the number of molecules used. The graph in Figure 1 shows the case for *k* = 37 and λ*_f_* = 0, 1, 2 and 4, which corresponds to the case analyzed in Section 4. We see that for *p* = 0.10 and λ_*f*_ = 1 a sharp transition occurs when the number of molecules increases from 30 to 50. At 70 or more molecules the AMS algorithm never (out of 20 experiments) fails to find the correct map, whereas for 20 or less molecules it invariably fails to find the correct map. For *p* = 0.20 (Figure 2), the transition (from probability of near 0 to near 1) occurs at a lower value of around 20–30 molecules. Compare this with the theoretical bounds on the number of molecules required from section 4 of between 30 and 100 (lower bound and upper bound respectively).

**Figure 1:**
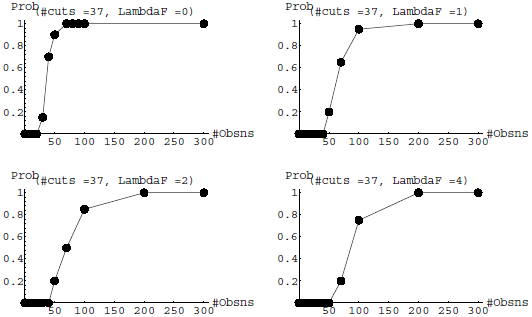
**Experimental Results:** #Cuts, *k* = 37, *σ* = 1.5bp, *p* = 0.1

**Figure 2:**
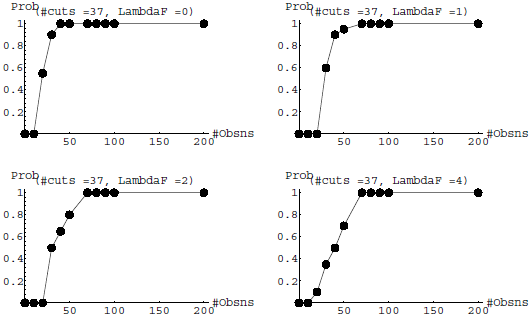
**Experimental Results:** #Cuts, *k* = 37, *σ* = 1.5bp, *p* = 0.2

When the number of (true) cuts in the molecules is changed to *k* = 20, 10, 5 and 2, similar graphs are obtained: Figures 4 and 5 show the results for the case *k* = 20; Figures 6 and 7, for the case *k* = 10; Figures 8 and 9, for the case *k* = 5; Figures 10 and 11, for the case *k* = 2; Figures 13 and 14, for the case *k* = 1 and Figure 15, for *k* = 0. The main trend is an increase in the number of molecules required as *k* is reduced down to *k* = 2: for instance, with *k* = 2 and *p* = 0.1, 500 molecules are required to find the correct map in every case (λ_*f*_ = 0, 1, 2 and 4), in contrast to just 200 for *k* = 37. This observation agrees with the theory from sections 4 and 5 which shows that the bounds increase slowly as *k* is decreased. However, the case *k* = 1, Figures 13 and 14, show that fewer molecules are required: e.g., with *p* = 0.1 and λ_*f*_ = 1, 200 molecules are sufficient to find the correct map. The reason is that orientation is less of a problem with only 1 cut.

**Figure 3:**
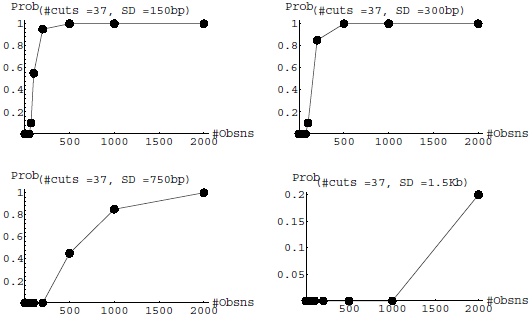
**Experimental Results:** #Cuts, *k* = 37, λ*_f_* = 1, *p* = 0.1

**Figure 4:**
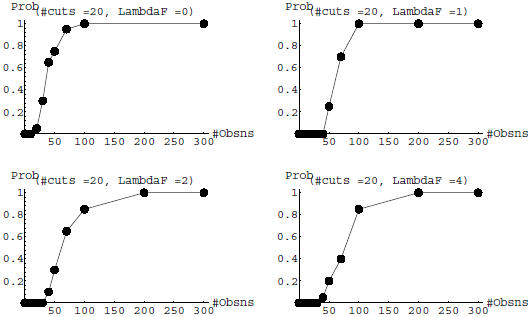
**Experimental Results:** #Cuts, *k* = 20, *σ* = 1.5bp, *p* = 0.1

**Figure 5:**
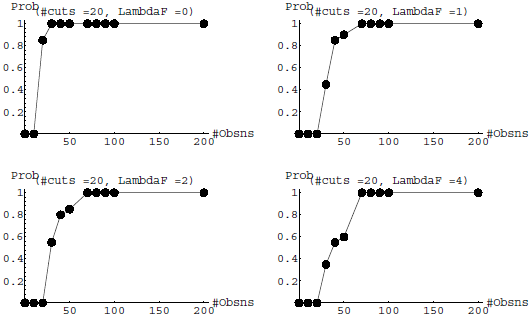
**Experimental Results:** #Cuts, *k* = 20, *σ* = 1.5bp, *p* = 0.2

**Figure 6:**
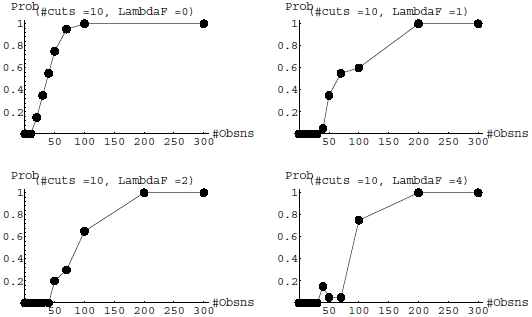
**Experimental Results:** #Cuts, *k* = 10, *σ* = 1.5bp, *p* = 0.1

**Figure 7:**
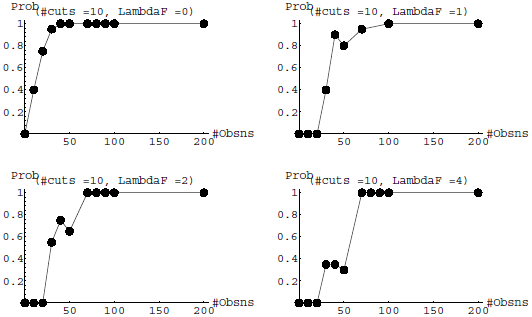
**Experimental Results:** #Cuts, *k* = 10, *σ* = 1.5bp, *p* = 0.2

**Figure 8:**
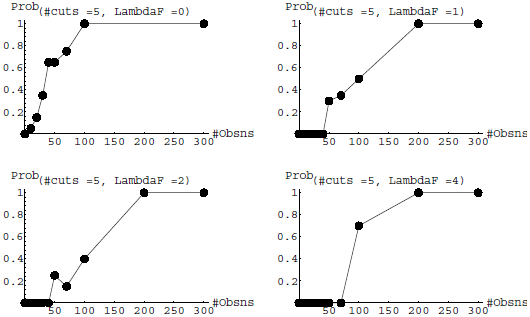
**Experimental Results:** #Cuts, *k* = 5, *σ* = 1.5bp, *p* = 0.1

**Figure 9:**
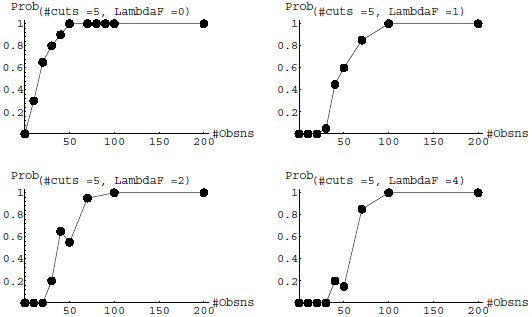
**Experimental Results:** #Cuts, *k* = 5, *σ* = 1.5bp, *p* = 0.2

**Figure 10:**
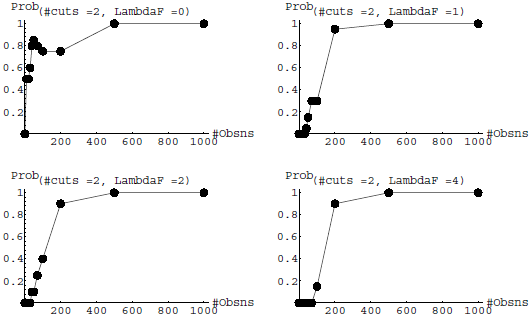
**Experimental Results:** #Cuts, *k* = 2, *σ* = 1.5bp, *p* = 0.1

**Figure 11:**
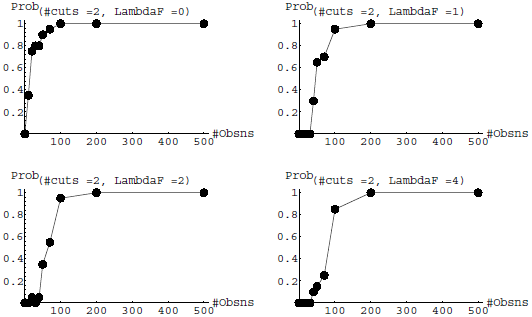
**Experimental Results:** #Cuts, *k* = 2, *σ* = 1.5bp, *p* = 0.2

**Figure 12:**
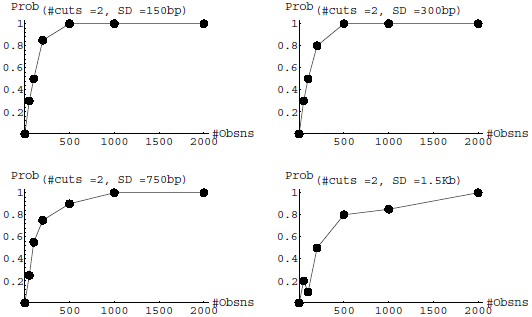
**Experimental Results:** #Cuts, *k* = 2, λ_*f*_ = 1, *p* = 0.1

**Figure 13:**
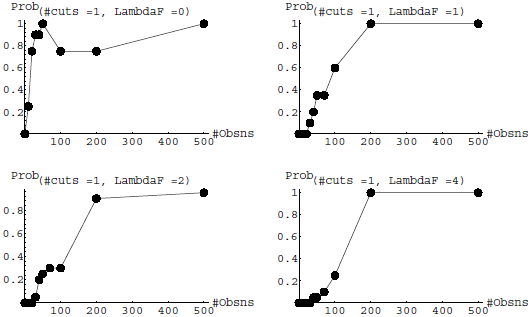
**Experimental Results:** #Cuts, *k* = 1, *σ* = 1.5bp, *p* = 0.1

**Figure 14:**
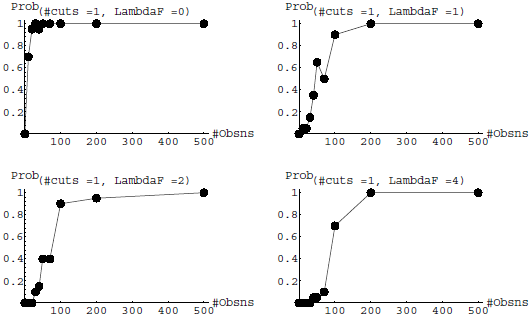
**Experimental Results:** #Cuts, *k* = 1, *σ* = 1.5bp, *p* = 0.2

**Figure 15:**
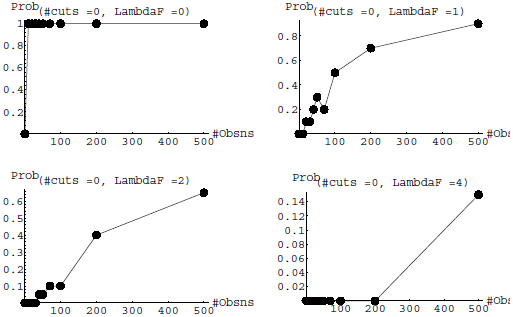
**Experimental Results:** #Cuts, *k* = 0

Figure 3 shows what happens with *k* = 37 when the sizing error is increased to 150bp, 300bp, 750bp and 1.5Kb, respectively. With *p* = 0.10 and λ_*f*_ = 1, the number of molecules required to find the correct map in every case increases from 200 to about 5000 as the sizing error increases. Figure 12 shows what happens at *k* = 2 when sizing error is increased similarly. In this case the number of molecules increases from an already larger value, but more slowly: it increases from 500 to 2000. While we do not have any theoretical bounds for this case, the intuition is that while it is harder to get the correct orientation with *k* = 2 than with *k* = 37, it is less likely that neighboring cuts will be confused with each other due to sizing errors when *k* = 2 than when *k* = 37.

## 8 Discussion: Genomic Mapping

The strategies for genome-wide genotype or haplotype mapping using single molecule optical maps are similar in spirit to the approaches for clone mapping, described in this paper; but there are also several differences in the details of the implementation as new and powerful heuristics need to be incorporated in order to tame the computational complexity of searching over the hypotheses space. The details of the algorithm can be found elsewhere [AnMS99]. We summarize below a 0-1 law applicable to the experiment design in this genomic-mapping setting; the derivation of the following result is in [AM01].

Consider an optical mapping experiment for genome-wide shotgun mapping for a genome of size *G* and involving *M* molecules each of length *L_d_*. Thus the coverage is *ML_d_*/*G*. Let the a fragment of true size *X* have a measured size ∼ 𝒩(*X*, *σ*^2^*X*). Let the average true fragment size be *L*, and the digestion rate of the restriction enzyme be *p*. Thus the average relative sizing error 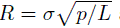 and the average size of aligned fragments will be *L*/*p*^2^. As usual, let *θ* represent the minimum “overlap threshold.” Hence the expected number of aligned fragments in a valid overlap is at least *n* = *θL*_*d*_*p*^2^/*L*. Let *d* = 1/*p*, the inverse of the digest rate. Feasible experimental parameters are those that result in an acceptable (e.g. ≤ 10^−3^) False Positive rate *FPT*:

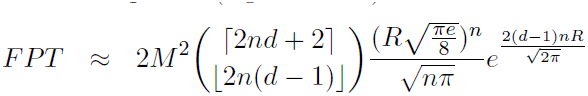

To achieve acceptable false positive rate, one needs to choose an acceptable value
for the experimental parameters: *p*, *σ*, *L_d_* and coverage. *FPT* exhibits a sharp phase transition in the space of experimental parameters. Thus the success of a mapping project depends extremely critically on a prudent combination of experimental errors (digestion rate, sizing), sample size (molecule length and number of molecules) and problem size (genome length). Relative sizing error can be lowered simply by increasing *L* with a choice of rarer-cutting enzyme and digestion rate can be improved by better chemistry.

As an example, for a human genome of size *G* = 3, 300Mb and a desired coverage of 6×, consider the following experiment. Assume a typical value of molecule length *L_d_* = 2*Mb*. If the enzyme of choice is pac I, the average true fragment length is about 25*Kb*. Assume a minimum overlap^1^ of *θ* = 30%. Assume that the sizing error for a fragment of 30*kb* is about 3.0*kb*, and hence *σ*2 = 0.3kb. With a digest rate of *p* = 82% we get an unacceptable FPT ≈ 0.0362. However just increasing p to 86% results in an acceptable *FPT* ≈ 0.0009. Alternately, reducing average sizing error from 3.0*kb* to 2.4*kb* while keeping *p* = 82% also produces an acceptable *FPT* ≈ 0.0007.

## Footnotes

1 This value should be selected to minimize *FPT*.

## Acknowledgment

Our thanks go to Naomi Silver, Rohit Parikh, Raghu Varadhan, Joel Spencer, Alan Frieze, Sylvain Cappel, Bruce Donald, Mike Wigler and Laxmi Parida for many helpful comments and encouragement.

## A1. Bound for section 4.2

Assume the same notation as in section §4.2: We can provide a somewhat better bound when *p* is small, i.e., *p* ≈ 1/*k*.

Let *α* be a function of *p* and *k*:

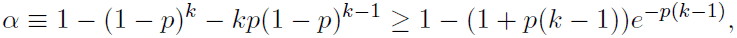

and let

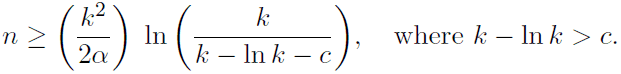

Construct a random subgraph *G_R_* = (*V*,*E_R_*) of *G* as follows: For any given observation with two or more cuts choose one edge at random from all the possible edges that the observation contributes to *G*. Discard those observations with fewer than two cuts. Thus with every observation, when we add an edge we do so uniformly randomly and independent of all the other edges chosen in *G_R_*. Note that the probability that an observation has two or more cuts is *α* and the probability that an edge is added to *G_R_* in an observation is 2*α*/*k*(*k* − 1) > 2*α*/*k*^2^.

**Table 1:** Summary of Experimental Results. Number of molecules necessary as functions of the parameters: #Cuts, *k* ∈ {0..37}, Digest rate *p* ∈ {0.1, 0.2} and λ_*f*_ ∈ {0, 1, 2, 4}.

| #Cuts, $k$ | Digest rate, $p_c$ | $\lambda_f = 0$ | $\lambda_f = 1$ | $\lambda_f = 2$ | $\lambda_f = 4$ |
| --- | --- | --- | --- | --- | --- |
| 37 | 0.1 | 50 | 100 | 200 | 200 |
|  | 0.2 | 40 | 70 | 70 | 70 |
| 20 | 0.1 | 100 | 100 | 200 | 200 |
|  | 0.2 | 30 | 70 | 70 | 70 |
| 10 | 0.1 | 100 | 200 | 200 | 200 |
|  | 0.2 | 50 | 70 | 70 | 70 |
| 5 | 0.1 | 100 | 200 | 200 | 200 |
|  | 0.2 | 50 | 100 | 100 | 100 |
| 2 | 0.1 | 500 | 500 | 500 | 500 |
|  | 0.2 | 100 | 200 | 200 | 200 |
| 1 | 0.1 | 70 | 200 | 200 | 200 |
|  | 0.2 | 30 | 200 | 200 | 200 |
| 0 | — | 1 | 500 | — | — |

For any pair [*f*(*h_i_*), *f*(*h_j_*)], the probability that this edge does not occur in *G_R_* is less than

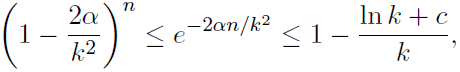

and thus the “edge-probability,” *p*_*e*_ (the probability that this edge occurs in *G_R_*) is

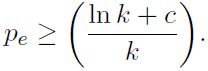

Thus by the well-known result on the connectivity in random graphs [Spe87], we see that with 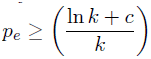,

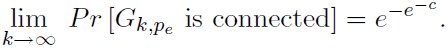

Note that

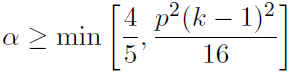

and if *k* ≫ *c* then

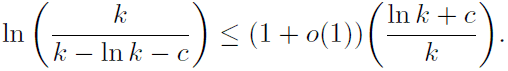

Thus it suffices for our purpose to choose

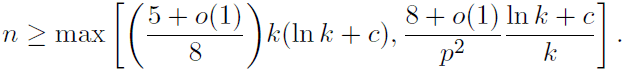

**Theorem 8.1** *Let *ϵ* be a positive constant and c* ≥ ln(2/*ϵ*). *Then for*

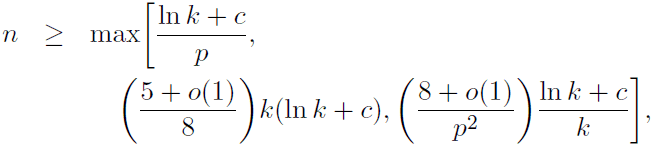

*with a probability at least* 1 − *ϵ*, *the correct ordered restriction map can be computed in O*(*nk*^2^) *time.*

Furthermore, we conclude that, for *p* ≥ 1/*k*, *n* = *O*(*k* log *k*) observations suffice to find the true map without any other prior knowledge of *p*.

## A2. Discretization

As before, let us assume that the clone DNA is of length *L* bps. Let ∆ represent a small subinterval and *δ* = ∆/*L*. Thus the unit length is partitioned into *M* = 1/δ = *L*/∆ consecutive subintervals. We write *r* = λ_*f*_ δ = ∆λ_*f*_/*L* to denote the probability that we shall observe one spurious cut in a subinterval.

Typical values for various clones may be as follows: for lambdas, *M* can range from 200 to 2, 000 and r ≈ 10^−3^–10^−4;^ for cosmids, *M* is 2,000–4,000 and *r* ≈ 10^−4^; for BACs *M* ≈ 15,000 and *r* ≈ 10^−4^. In general, even for significantly smaller (but still realistic) values of *M*, *r* ≪ *p*.

**Bounds**

We write *p̂* = *p* + *r* − *pr* to denote the probability that a subinterval contains a true or spurious cut site. We will use the following simplifying assumption:

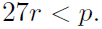

More precisely, *p̂*/6*r* > 2*e* − 1.

We summarize the bounds as follows:

Theorem 8.2 *Let *ϵ* be a positive constant and c* ≥ ln(5/*ϵ*). *Then for*

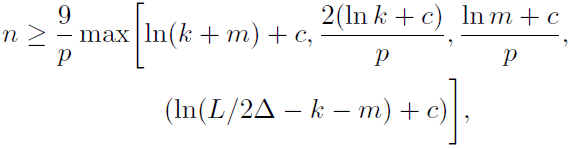

(*L* > 2∆ and *r* < *p*/27), *the probability that the correct ordered restriction map can be computed in O*(*n*(*L* + *k*^2^ + *m*)) time is at least 1 − *ϵ*.

We will now introduce two parameters *ϵ*_1_ = *p̂*/6*r* and *ϵ*_0_, and guarantee that *ϵ*_1_ > 2*e* − 1 and *ϵ*_0_ ≥ 1/2. Furthermore, we have

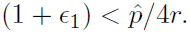

**Phase 1 a**

In phase 1 a, our goal is to construct the set

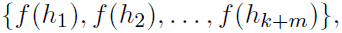

by considering the observation-based sets

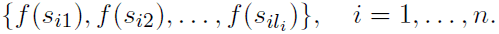

and including only those *f*(*s_ij_*)’s that occur in *significantly large numbers* of times, determined by a threshold *Th_1_*. Suppose that a location *f*(*h*) corresponds to a true location, then the number of *f*(*s_ij_*)’s equal to *f*(*h*) must follow a Binomial distribution ∼ *S*(*n*, *p̂*), if it is an asymmetric cut and ∼ *S*(n, 2*p̂*), if it is a symmetric cut. If on the other hand, *f*(*h*) does not correspond to any true location, then the number of *f*(*s_ij_*)’s equal to this *f*(*h*) must follow a Binomial distribution ∼ *S*(*n*, 2*r*).

If we set the threshold at

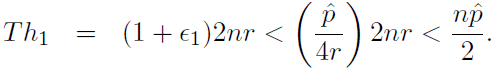

then *Th*_1_ = (1 − *ϵ*_0_)*n*p̂**, where *ϵ*_0_ ≥ 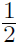.

By assumption (statement of the theorem above),

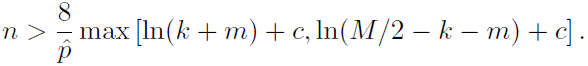

Thus (using the Chernoff bound [ASE92])

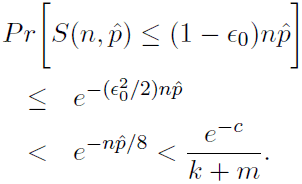

Thus the probability that all the correct cuts appear in the computed set is bounded from below by *e*^−*e*^−*c*^^.

Again, using the Chernoff bound [ASE92] in the other direction, we get

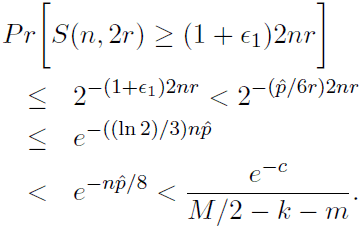

Thus the probability that no spurious cut appears in the computed set < 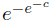.

**Phase 1 b**

In phase 1 b, our goal is to construct the set of asymmetric cuts

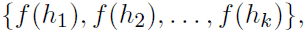

by eliminating the symmetric cuts. Suppose that a location *f*(*h*) corresponds to a symmetric true cut site, then the number of times an observation has sites at *s*′ = *f*(*h*) and *s*′′ = *f*(*h*)^*R*^ must follow a Binomial distribution ∼ *S*(*n*, *p̂*^2^). If on the other hand, *f*(*h*) is not a symmetric site, then the corresponding number must follow a Binomial distribution ∼ *S*(*n*, *p̂r*).

If we set the threshold at

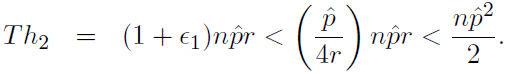

then *Th*_2_ = (1 − *ϵ*_0_)n*p̂*2, where *ϵ*_0_ ≥ 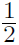.

By assumption (statement of the theorem above),

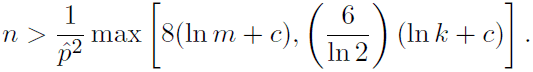

Thus (using the Chernoff bound [ASE92])

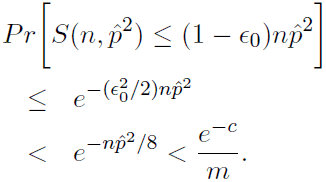

Thus the probability that all the symmetric cuts are correctly classified is bounded from below by *e*^−*e*−*c*^.

Again, using the Chernoff bound [ASE92] in the other direction, we get

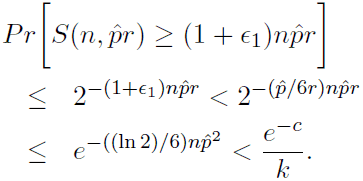

Thus the probability that no symmetric cut is misclassified is bounded from below by 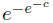.

**Phase 2**

The proof proceeds in a manner similar to the one given for the non-discretized case. In phase 2, our goal is to assign consistent sign labels to the asymmetric cuts

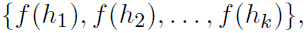

so that the final map can be constructed correctly with high probability.

Let *S*_*h*1_ denote the set of observations containing a cut site matching *f*(*h*_1_), and 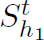 denote the set containing a true cut site matching *f*(*h*_1_). Note that 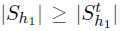 and 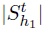 follows a Binomial distribution ∼ *S*(*n*, *p*). Using the Chernoff bound, we have

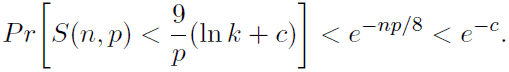

Let *n*_1_ be the number cut sites matching *h*_1_. Thus

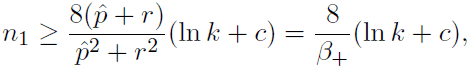

with a probability > 1 −*e*^−*c*^. Here *β*_+_ ≡ (*p̂*^2^ + *r*^2^)/(*p̂* + *r*).

Consider a potential edge [*f*(*h*_1_), *f*(*h_i_*)]. Let ni denote the number of times two cut sites matching *f*(*h*_1_) and *f*(*h_i_*), respectively, appear in the same half [either in (0, 1/2) or in (1/2, 1)] in an observation in S_h1_. If the correct edge labeling is +1 then n_i_ has a Binomial distribution ∼ *S*(*n*_1_, *β*_+_), where *β*_+_ ≡ (*p̂*^2^ + *r*^2^)/(*p̂* + *r*). If on the other hand, the correct edge labeling is −1 then ni has a Binomial distribution ∼ *S*(*n*_1_, *β*_−_), where *β*_−_ ≡ (2*p̂r*)/(*p̂* + *r*)).

Set the threshold at

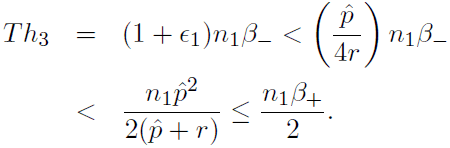

Thus *Th*_3_ = (1 − *ϵ*_0_)*n*_1_*β*_+_, where *ϵ*_0_ ≥ 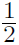.

Thus (using the Chernoff bound [ASE92])

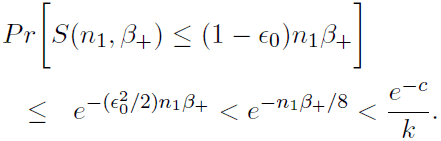

Again, using the Chernoff bound [ASE92] in the other direction, we get

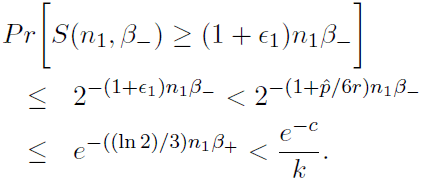

Thus it follows that the probability that all the edges receive the correct edge labeling is > (1 − *e*^−*c*^)*e*^−*e*^−*c*^^. This concludes the proof.

## References

[ASE92] N. Alon, J.H. Spencer and P. Erdös (1992). The Probabilistic Method. Wiley Interscience.(John Wiley & Sons, Inc., New York.)

[AM01] T.S. Anantharaman and B. Mishra (2001). A Probabilistic Analysis of False Positives in Optical Map Alignment and Validation. Algorithms in Bioinformatics, First International Workshop, WABI 2001 Proceedings **LNCS 2149**, 27–40. (Springer-Verlag, New York.)

[AMS97] T.S. Anantharaman, B. Mishra and D.C. Schwartz (1997). Genomics via Optical Mapping II: Ordered Restriction Maps. Journal of Computational Biology 4(2), 91–118.

[Ana+97b] T.S. Anantharaman et al. (1997). Statistical Algorithms for Optical Mapping of the Human Genome. 1997 Genome Mapping and Sequencing Conference, Cold Spring Harbor, New York.

[AnMS99] T.S. Anantharaman, B. Mishra and D.C. Schwartz (1999). Genomics via Optical Mapping III: Contiging Genomic DNA and Variations. Proceedings 7th Intl. Cnf. on Intelligent Systems for Molecular Biology: ISMB ’99 7, 18–27. (AAAI Press, New York).

[AsMS99] C. Aston, B. Mishra and D.C. Schwartz (1999). Optical Mapping and Its Potential for Large-Scale Sequencing Projects. Trends in Biotechnology 17, 297–302.

[Cai+98] W. Cai et al., High Resolution Restriction Maps of Bacterial Artificial Chromosomes Constructed by Optical Mapping,” Proc. National Academy of Science, (In Press), 1998.

[Giacalone+00] J. Giacalone et al. (2000). Optical Mapping of BAC Clones from the Human Y Chromosome DAZ Locus. Genome Research 10(9), 1421–1429.

[Jing+99] J. Jing et al., “Optical Mapping of *Plasmodium falciparum* Chromosome 2,” Genome Research, 9:175–181, 1999.

[KS98] R. Karp and R. Shamir (2000). Algorithms for Optical Mapping. Journal of Computational Biology 7(1–2), 303–316.

[Lai+99] Z. Lai et al. (1999). A Shotgun Sequence-Ready Optical Map of the Whole *Plasmodium falciparum* Genome. Nature Genetics 23(3), 309–313.

[Lim+01] A. Lim et al. (2001). Shotgun Optical Maps of the Whole *Escherichia coli* 0157:H7 Genome. Genome Research 11(9), 1584–1593.

[lin+99] J. Lin et al. (1999). Whole Genome Shotgun Optical Mapping of *Deinococcus radiodurans*. Science 285(5433), 1558–1562.

[Mishra03] B. Mishra (2003). Optical Mapping. Encyclopedia of the Human Genome, Nature Publishing Group. (Macmillan Publishers Limited, London.)

[MP00] B. Mishra and L. Parida (2000). Partitioning Single-Molecule Maps into Multiple Populations: Algorithms And Probabilistic Analysis. Discrete Applied Mathematics (The Computational Molecular Biology Series) 104(1–3), 203–227.

[MP96] S. Muthukrishnan and L. Parida (1997). Towards Constructing Physical Maps by Optical Mapping: An Effective Simple Combinatorial Approach. In Proceedings First Annual Conference on Computational Molecular Biology, (RECOMB ’97), ACM Press, 209–215.

[Parida98] L. Parida (1998). Algorithmic Techniques in Computational Genomics, PhD Thesis, Computer Science Department, New York University, New York.

[Sam+95] A. Samad et al. (1995). Mapping the Genome One Molecule At a Time—Optical Mapping. Nature 378, 516–517.

[Skiadas+99] J. Skiadas et al. (1999). Optical PCR: Genomic Analysis by Long-Range PCR and Optical Mapping. Mammalian Genome 10, 1005–1009.

[Spe87] J. Spencer (1987). Ten Lectures on the Probabilistic Method. (Society for Industrial and Applied Mathematics, Philadelphia, PA.)

[Zhou+02] S. Zhou et al. (2002). A Whole-Genome Shotgun Optical Map of *Yersinia pestis* Strain KIM. Appl Environ Microbiol 68(12), 6321–6331.

